# Sagittal abdominal diameter and waist circumference are equally good as identifiers of cardiometabolic risk

**DOI:** 10.1101/598078

**Authors:** Grith Møller, Christian Ritz, Louise Kjølbæk, Stine Vuholm, Sanne Kellebjerg Korndal, Thomas Meinert Larsen, Oluf Pedersen, Wim Saris, Arne Astrup, Lotte Lauritzen, Mette Kristensen, Mads Vendelbo Lind

## Abstract

**Background:** Body mass index (BMI) and waist circumference (WC) are commonly used markers of cardiometabolic risk. However, sagittal abdominal diameter (SAD) has been proposed to be a better marker of intra-abdominal obesity compared to WC and might better associate with metabolic disturbances in high-risk populations. The objective of this study was to compare SAD, WC, and BMI as determinants of an adverse metabolic phenotype.

**Method:** Anthropometric and metabolic measures of 1516 overweight or obese individuals with features of the metabolic syndrome were included to examine differences between SAD, WC and BMI as measures of an adverse metabolic phenotype. Multiple linear regression and logistic regression models were used to investigate the association between SAD, WC, and BMI and markers of metabolic syndrome, insulin resistance, blood lipids, and low grade inflammation.

**Results:** Both SAD and WC correlated with BMI, but as BMI increased, SAD proportionately estimated higher abdominal adiposity compared to WC (slope = 0.0037 (0.0029; 0.0046), p<0.0001). We did not find major differences between SAD, WC and BMI in explained variance in models with the different markers of metabolic risk. Furthermore, we did not find differences between SAD and WC in the ability to identify individuals with metabolic syndrome according to the International Diabetes Federation (IDF) cut-offs, but a few differences from BMI were indicated but mostly before adjustments. Moreover, the differences between SAD and WC associations were not modified by sex or degree of adiposity, but identification of individuals with a metabolic phenotype was generally better in women.

**Conclusion:** These data indicate that SAD and WC are equally good indicators of an adverse metabolic phenotype. Thus, from a public health perspective choice of anthropometric measure may depend only on what is the most practical method in a given situation.

## Introduction

Obesity is one of the most important risk factors for metabolic syndrome, cardiovascular disease (CVD) and type 2-diabetes (T2D). Body mass index (BMI) is often used to evaluate overweight and obesity, but BMI does not distinguish between lean and fat mass. Furthermore, BMI does not provide information on fat distribution, and therefore does not specifically reflect central obesity which is associated with an adverse metabolic phenotype and essential for the association between obesity and lifestyle disease (1).

When assessing abdominal obesity it is important to distinguish between subcutaneous adipose tissue (SAT) and visceral adipose tissue (VAT) as VAT is the most metabolically adverse type of adipose tissue (1). Simple anthropometric indicators such as waist circumference (WC) and sagittal abdominal diameter (SAD) are used to examine abdominal obesity, and have both been shown to be better indicators measures of an adverse metabolic phenotype compared to BMI alone (2). WC is a commonly used measure of abdominal obesity, but SAD has been proposed to be a better marker for VAT and thus might better have improved predictive ability of metabolic disturbances (3,4).

SAD is measured as the height of the abdomen in a supine position, which increase with VAT accumulation as VAT is contained within the abdominal cavity, while SAT in the supine position will be distributed sideways, due to the force of gravity (5,6). Adjusting measures of WC and SAD for height (as in BMI) have been shown to improve prediction of cardiometabolic risk and thus is suggested to also improve the detection of adverse metabolic phenotypes (7).

Some studies have suggested that sex and degree of obesity might affect the predictive ability of SAD and WC measurements differently (8,9). One study found that SAD compared to WC was a stronger risk marker in women compared to men (8). This could be due to the sex specific fat distribution as women have a higher ratio of SAT to VAT mass, which can lead to formation of a so-called “abdominal apron” and make measurements of WC particularly tricky. Similarly, difficulties might also make SAD a better measure than WC at higher levels of adiposity.

The overall aim of this study was to investigate whether SAD, WC and BMI associate differently with metabolic syndrome and cardiometabolic risk, compromising markers of insulin resistance, dyslipidemia, low grade inflammation and blood pressure. We hypothesized that SAD is stronger associated with cardiometabolic health and a better measurement for identifying individuals with metabolic syndrome markers above established cut-offs compared to WC and BMI. Furthermore, we hypothesise that height-adjustment of SAD and WC improves the association with adverse metabolic syndrome markers. Finally, we hypothesise that these associations depend on sex and that a stronger association for SAD compared to WC is most pronounced at higher BMIs (>35 kg/m^2^). These hypotheses were examined with data from several cohorts of primarily overweight and obese individuals with an adverse metabolic phenotype.

## Methods

### Study design

This study included cross-sectional baseline data from six human intervention trials: 3G, OPUS, DIOGENES, RIGHT, MyNewGut (MNG), and PROKA (10–15). The studies were registered at http://www.clinicaltrials.gov; 3G (NCT01719913 and NCT01731366); SHOPUS (NCT01195610); DIOGENES (NCT00390637); RIGHT (NCT02358122); MNG: (NCT02215343); PROKA (NCT01561131) and approved by the Research Ethics Committees of the Capital Region of Denmark in accordance with Helsinki Declaration 3G (H-2-2012-064 and H-2-2012-065); SHOPUS (H-3-2010-058); RIGHT (H-1-2014-062); MNG (H-4-2014-052); and PROKA (H-2-2011-145) or the local ethical committees in the respective countries (16).

### Participants

Individuals included in the present study, were primarily overweight or obese (98%) at risk of the metabolic syndrome. We used the International Diabetes Federation (IDF) metabolic syndrome definitions (17). For impaired fasting glucose we used both the 5.6 mmol/L IDF cut-off and a 6.1 mmol/L cut-off.

### Anthropometric, laboratory and analytical procedures

Body weight, height, WC and SAD were measured by standard anthropometric procedures. SAD was measured twice to the closest 0.1 cm using a Holtain-Kahn Abdominal Caliper (Holtain Ltd, Crymych, United Kingdom) and we used the mean of the two measurements. The subjects were in supine position when measured and were asked to bend their knees to a 45 degree angle and keep their feet flat on the examination table. SAD was measured as the distance between the highest point of the abdomen and the back as assessed by the distance between the two blades of the caliper (by using a spirit level). The measurement was made at the end of a normal exhalation. WC was measured between the bottom of the ribs and the top of the hipbone. WC was measured twice to the nearest 0.5 cm using a non-elastic flexible measuring tape and the mean of the two measurements was used. All cohorts used a similar standard operating procedure for the anthropometric measurements.

Blood samples for all cohorts were drawn after overnight fasting. Standard clinical measurements of plasma glucose, insulin, total-, LDL- and HDL cholesterol, triglycerides, C-reactive protein (CRP), free fatty acids (FFA), and adiponectin were performed as well as measurements of serum inflammatory markers interleukin (IL)-6 and tumor necrosis factor (TNF)-α (Supplemental methods section). Blood pressure was measured using a digital blood pressure monitor (Supplemental methods section).

### Statistical analysis

The statistical analyses were performed using R (v.3.5.1, R Core Team, Vienna, Austria) (18). Only baseline data from the studies were used in the analyses. Pearson correlation was used for evaluating the association between anthropometric measures. Simple linear regression of mean-centered data was used to evaluate any bias between measurements of SAD and WC across the BMI range, and reported as the slope. Multiple linear regression models were used to investigate the association between SAD, WC, and BMI and markers of metabolic syndrome, insulin resistance, blood lipids and measurements of low grade inflammation. Unadjusted analyses as well as analyses adjusted for age, sex, smoking and study were conducted. Similar analyses were done for the SAD-to-height and WC-to-height ratios. The ability of SAD, WC, and BMI to identify individuals with metabolic syndrome markers above the IDF cut-offs was evaluated by means of area under the receiver operating characteristics (ROC) curve using logistic regression models. Unadjusted analyses as well as analyses adjusted for age, sex, smoking and study were conducted

## Results

The six cohorts provided a total of 1516 individuals (485 men and 1031 women) (Table 1). The participants had an average age of 42 years, and their BMI ranged from 21 to 52 kg/ m^2^, WC ranged from 64-155 cm and SAD ranged from 15-39 cm.

**Table 1.** Characteristics of the participants at baseline (n = 1516)

| Variables | n | 3G | n | SHOPUS | n | DIOGENES | n | RIGHT | n | MNG | n | PROKA | n | Across studies |
| --- | --- | --- | --- | --- | --- | --- | --- | --- | --- | --- | --- | --- | --- | --- |
| Age (years) | 117 | 48.70 ± 11.25<sup>a</sup> | 181 | 42.10 ± 13.08<sup>b</sup> | 902 | 41.14 ± 6.26<sup>b,c</sup> | 73 | 50.85 ± 9.26<sup>c</sup> | 29 | 43.97 ± 11.53<sup>a,b</sup> | 214 | 40.03 ± 10.68<sup>b</sup> | 1516 | 42.20 ± 9.17 |
| Men % (M/F) | 117 | 39.3 (46/71)<sup>a</sup> | 181 | 31.0 (53/128)<sup>a</sup> | 902 | 33.4 (301/601)<sup>a,b</sup> | 73 | 43.8 (32/41)<sup>b</sup> | 29 | 28.0 (8/21)<sup>a</sup> | 214 | 39.5 (45/169)<sup>a</sup> | 1516 | 32.0 (485/1031) |
| Weight (kg) | 117 | 85.9 ± 13.0<sup>a</sup> | 181 | 89.9 ± 17.1<sup>a</sup> | 902 | 99.6 ± 17.4<sup>b</sup> | 73 | 83.5 ± 9.1<sup>a</sup> | 29 | 88.0 ± 13.7<sup>a</sup> | 214 | 96.6 ± 13.6<sup>b</sup> | 1516 | 96.0 ± 17.0 |
| Sagittal abdominal diameter (cm) | 117 | 22.9 ± 2.9<sup>c,d</sup> | 181 | 23.0 ± 3.2<sup>c</sup> | 902 | 25.2 ± 3.8<sup>a</sup> | 73 | 21.5 ± 2.1<sup>d</sup> | 29 | 22.0 ± 2.5<sup>c,d</sup> | 214 | 24.0 ± 2.5<sup>b</sup> | 1516 | 24.4 ± 3.6 |
| Waist circumference (cm) | 117 | 100.0 ± 9.0<sup>b,c</sup> | 181 | 100.1 ± 12.4<sup>b,c</sup> | 902 | 107.5 ± 13.0<sup>a</sup> | 73 | 96.0 ± 8.0<sup>c</sup> | 29 | 96.5 ± 8.8<sup>c</sup> | 214 | 103.2 ± 10.6<sup>b</sup> | 1516 | 104.70 ± 12.60 |
| Body-mass index (kg/m<sup>2</sup>) | 117 | 28.7 ± 3.5<sup>a</sup> | 181 | 30.3 ± 4.9<sup>c</sup> | 902 | 34.5 ± 4.8<sup>b</sup> | 73 | 27.8 ± 1.9<sup>a</sup> | 29 | 30.1 ± 3.1<sup>a,c</sup> | 214 | 33.2 ± 3.3<sup>d</sup> | 1516 | 33.0 ± 5.0 |
| Smoking, % (no/yes) | 117 | 7 (109/8)<sup>a</sup> | 181 | 22 (136/39)<sup>b</sup> | 851 | 28 (616/235)<sup>b</sup> | 73 | 0 (73/0)<sup>b</sup> | 29 | 0 (29/0)<sup>a</sup> | 214 | 0 (214/0)<sup>c</sup> | 1459 | 19 (1177/282) |
| Fasting insulin (pmol/L) | 114 | 65.0 ± 32.1<sup>a</sup> | 181 | 75.4 ± 45.3 | 799 | 83.4 ± 72.1<sup>b</sup> | 73 | 74.2 ± 39.4 | 28 | 43.6 ± 30.3<sup>a</sup> | 210 | 74.6 ± 44.5 | 1405 | 78.3 ± 61.2 |
| Fasting glucose (mmol/L) | 117 | 5.7 ± 0.6<sup>c</sup> | 181 | 5.3 ± 0.5<sup>b</sup> | 817 | 5.0 ± 0.7<sup>a</sup> | 73 | 5.9 ± 0.6<sup>c</sup> | 28 | 5.5 ± 0.4<sup>b</sup> | 210 | 5.7 ± 0.7<sup>c</sup> | 1426 | 5.3 ± 0.7 |
| 2-h glucose (mmol/L) |  | - | 181 | 5.6 ± 1.4<sup>a</sup> | 794 | 6.7 ± 2.2<sup>b</sup> |  | - |  | - |  | - | 975 | 6.5 ± 2.1 |
| HOMA-IR | 114 | 2.4 ± 1.3<sup>b,c</sup> | 181 | 2.5 ± 1.6<sup>c</sup> | 798 | 3.2 ± 2.9<sup>a</sup> | 73 | 2.8 ± 1.6<sup>a,c</sup> | 28 | 1.0 ± 0.7<sup>b</sup> | 210 | 1.9 ± 1.4<sup>b,c</sup> | 1404 | 2.8 ± 2.4 |
| Matsuda index |  | - | 180 | 5.7 ± 3.0<sup>a</sup> | 783 | 5.3 ± 3.3<sup>b</sup> |  | - |  | - |  | - | 963 | 5.4 ± 3.2 |
| TNF-α (pg/mL) | 117 | 1.8 ± 2.2<sup>a</sup> | 181 | 1.2 ± 2.2<sup>b</sup> | | - | | - | | - | | - | 298 | 1.4 ± 2.2 |
| IL-6 (pg/mL) | 116 | 1.7 ± 1.6<sup>a</sup> | 180 | 1.3 ± 0.9<sup>b</sup> |  | - | 73 | 2.1 ± 1.6<sup>c</sup> |  | - |  | - | 369 | 1.6 ± 1.3 |
| CRP (mg/L) | 117 | 1.5 ± 1.8<sup>a</sup> | 181 | 2.6 ± 2.3<sup>c</sup> | 744 | 3.3 ± 2.3<sup>d</sup> | 72 | 1.8 ± 1.6<sup>a,c</sup> | 26 | 2.0 ± 2.0<sup>a,c</sup> |  | - | 1113 | 2.9 ± 2.3 |
| Adiponectin (μg/mL) | | - | | - | 832 | 9.1 ± 4.3 | | - | | - | | - | 832 | 9.1 ± 4.3 |
| Total cholesterol (mmol/L) | 117 | 5.3 ± 1.0<sup>c</sup> | 180 | 4.6 ± 0.9<sup>b</sup> | 832 | 4.9 ± 1.0<sup>a</sup> | 73 | 5.2 ± 0.9<sup>a,c</sup> | 28 | 5.0 ± 0.9 | 210 | 5.3 ± 0.9<sup>c</sup> | 1440 | 5.0 ± 1.0 |
| HDL cholesterol (mmol/L) | 117 | 1.3 ± 0.3<sup>b</sup> | 180 | 1.2 ± 0.3<sup>a</sup> | 834 | 1.2 ± 0.3<sup>a</sup> | 73 | 1.4 ± 0.3<sup>b</sup> | 28 | 1.4 ± 0.4<sup>b</sup> | 210 | 1.4 ± 0.4<sup>b</sup> | 1442 | 1.2 ± 0.3 |
| LDL cholesterol (mmol/L) | 117 | 3.1 ± 0.7 | 180 | 3.0 ± 0.8 | 828 | 3.1 ± 0.9 | 73 | 3.3 ± 0.8 | 28 | 3.1 ± 0.9 | 210 | 3.2 ± 0.8 | 1430 | 3.1 ± 0.8 |
| Triglycerides (mmol/L) | 117 | 1.3 ± 0.6<sup>b,c</sup> | 180 | 1.2 ± 0.6<sup>b</sup> | 822 | 1.4 ± 0.7<sup>c</sup> | 73 | 1.2 ± 0.6<sup>b</sup> | 28 | 2.8 ± 1.1<sup>a</sup> | 210 | 1.4 ± 0.8<sup>c</sup> | 1430 | 1.4 ± 0.7 |
| FFA (mmol/L) | 117 | 0.48 ± 0.16<sup>a</sup> | 159 | - | 735 | 0.65 ± 0.32<sup>b</sup> |  | - |  | - | 211 | 0.46 ± 0.16<sup>c</sup> | 1063 | 0.6 ± 03 |
| Systolic blood pressure (mmHg) | 117 | 129 ± 13<sup>a</sup> | 181 | 122 ± 14<sup>b</sup> | 885 | 125 ± 15<sup>a,b</sup> | 73 | 129 ± 17<sup>a</sup> | 29 | 120 ± 15<sup>b,c</sup> | 214 | 118 ± 11<sup>c</sup> | 1499 | 124.0 ± 14.5 |
| Diastolic blood pressure (mmHg) | 117 | 81 ± 9<sup>a</sup> | 181 | 81 ± 10<sup>a</sup> | 885 | 77 ± 11<sup>b</sup> | 73 | 85 ± 12<sup>a</sup> | 29 | 78 ± 10<sup>b,c</sup> | 214 | 76 ± 9<sup>c</sup> | 1499 | 78.3 ± 10.8 |

Data shown as mean ± SD. <sup>a,b,c</sup> Values with different superscripts letters indicate statistically significant differences between studies, assessed by analysis of variance or chi-square tests, p < .05. Abbreviations: CRP, C-reactive protein; FFA, Free fatty acids; HOMA-IR, Homeostatic model assessment for insulin resistance; IL-6, Interleukin 6, TNF- $\alpha$ , Tumor necrosis factor $\alpha$ .

The three anthropometric measures were highly correlated (r=0.82 for SAD and WC; r=0.74 for SAD and BMI; r=0.76 for WC and BMI). However, there was a systematic difference between the correlations for SAD and WC over the range of BMI range of 21-52 kg/m^2^. As BMI increased there was a significant trend for SAD to measure proportionately higher abdominal adiposity compared to WC. This was highly significant for men, women, and men and women combined (**Figure 1**).

**Figure 1.**
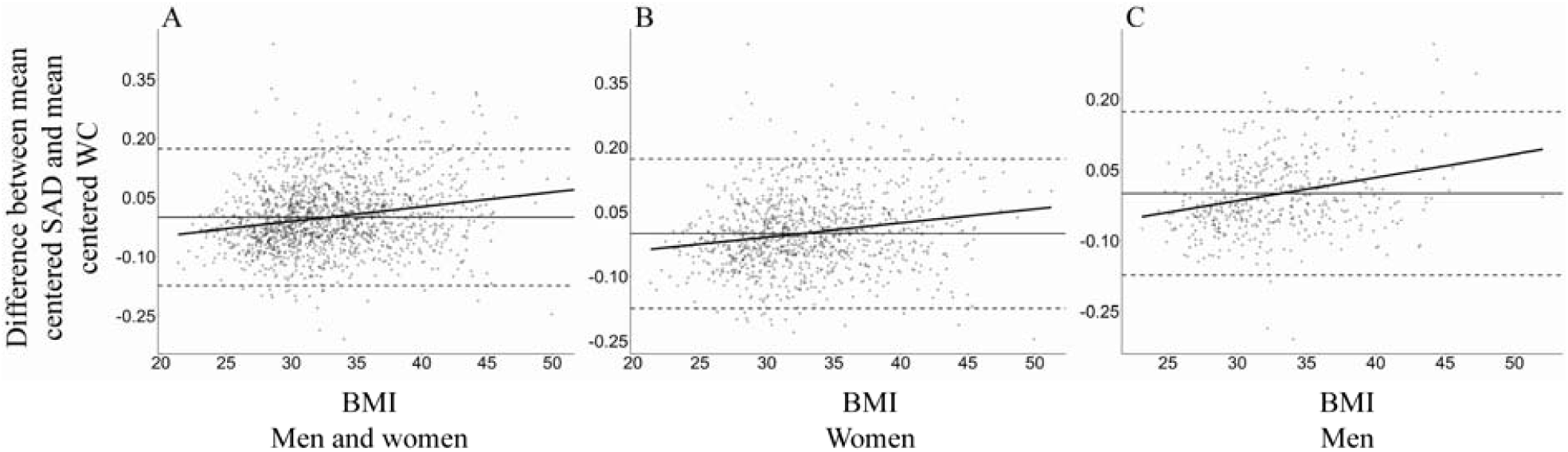
Difference between mean-centered sagittal abdominal diameter (SAD) and waist circumference (WC) across the body-mass index (BMI) range of the participants. A) Men and women combined slope = 0.0037 (0.0029; 0.0046), p<0.0001, n=1516. B). Women: slope = 0.0032 (0.0022; 0.0043), p<0.0001, n=1031. C). Men: slope = 0.0049 (0.0035; 0.0064), p<0.0001, n=485.

### Associations between anthropometric measures and cardiometabolic health

BMI, WC and SAD were associated with all markers of an adverse metabolic phenotype (Table 2). There were little difference between the measures in terms of R^2^ values between SAD and WC, indicating that they explained the variation in cardiometabolic measures equally well (Table 2). Furthermore, there were no differences between the variance explained by SAD-to-height ratio and WC-to-height ratio (Supplemental table 1) and the variation explained by these measures did not from those of SAD and WC alone.

**Table 2.** Associations between SAD, WC, BMI and markers of cardiometabolic health

|  | n | SAD (cm) |  |  | WC (cm) |  |  | BMI (kg/m2) |  |  |
| --- | --- | --- | --- | --- | --- | --- | --- | --- | --- | --- |
|  |  | Slope (CI) | p-value | R<sup>2</sup> | Slope (CI) | p-value | R<sup>2</sup> | Slope (CI) | p-value | R<sup>2</sup> |
| **Fasting insulin (pmol/L)** |  |  |  |  |  |  |  |  |  |  |
| Unadjusted | 1405 | 14.6 (12.7, 16.5) | <0.001 | 0.139 | 16.8 (14.9, 18.7) | <0.001 | 0.180 | 15.0 (11.1, 14.9) | <0.001 | 0.110 |
| Adjusted | 1354 | 14.7 (12.5, 16.9) | <0.001 | 0.173 | 17.7 (15.5, 19.9) | <0.001 | 0.213 | 14.8 (12.6, 17.0) | <0.001 | 0.173 |
| **Fasting glucose (mmol/L)** |  |  |  |  |  |  |  |  |  |  |
| Unadjusted | 1426 | 0.04 (0.007, 0.1) | 0.019 | 0.004 | 0.1 (0.1, 0.1) | <0.001 | 0.014 | -0.02 (-0.1, 0.01) | 0.180 | 0.001 |
| Adjusted | 1376 | 0.09 (0.06, 0.1) | <0.001 | 0.252 | 0.1 (0.1, 0.2) | <0.001 | 0.263 | 0.1 (0.1, 0.1) | <0.001 | 0.256 |
| **2-h glucose (mmol/L)** |  |  |  |  |  |  |  |  |  |  |
| Unadjusted | 975 | 0.3 (0.2, 0.4) | <0.001 | 0.023 | 0.2 (0.1, 0.3) | <0.001 | 0.016 | 0.3 (0.2, 0.4) | <0.001 | 0.026 |
| Adjusted | 925 | 0.3 (0.2, 0.4) | <0.001 | 0.083 | 0.2 (0.1, 0.4) | <0.001 | 0.077 | 0.2 (0.1, 0.4) | <0.001 | 0.077 |
| **HOMA-IR** |  |  |  |  |  |  |  |  |  |  |
| Unadjusted | 1404 | 0.6 (0.5, 0.7) | <0.001 | 0.151 | 0.7 (0.6, 0.8) | <0.001 | 0.198 | 0.5 (0.4, 0.6) | <0.001 | 0.110 |
| Adjusted | 1353 | 0.5 (0.5, 0.6) | <0.001 | 0.247 | 0.7 (0.6, 0.7) | <0.001 | 0.285 | 0.6 (0.5, 0.6) | <0.001 | 0.248 |
| **Matsuda index** |  |  |  |  |  |  |  |  |  |  |
| Unadjusted | 963 | -0.9 (1.1, -0.8) | <0.001 | 0.121 | -1.1 (-1.3, -1.0) | <0.001 | 0.181 | -0.8 (-0.9, -0.6) | <0.001 | 0.081 |
| Adjusted | 913 | -0.8 (-1.01, -0.7) | <0.001 | 0.140 | -1.1 (-1.3, -0.9) | <0.001 | 0.191 | -0.8 (-1.0, -0.6) | <0.001 | 0.137 |
| **TNF-α (pg/mL)** |  |  |  |  |  |  |  |  |  |  |
| Unadjusted | 298 | 0.1 (0.001, 0.2) | 0.047 | 0.013 | 0.1 (0.001, 0.2) | 0.048 | 0.013 | 0.1 (-0.02, 0.1) | 0.123 | 0.008 |
| Adjusted | 292 | 0.1 (-0.002, 0.2) | 0.043 | 0.221 | 0.1 (-0.002, 0.2) | 0.056 | 0.220 | 0.1 (0.04, 0.2) | 0.003 | 0.233 |
| **IL-6 (pg/mL)** |  |  |  |  |  |  |  |  |  |  |
| Unadjusted | 369 | 0.2 (0.1, 0.3) | <0.001 | 0.048 | 0.2 (0.1, 0.3) | <0.001 | 0.036 | 0.2 (0.1, 0.3) | <0.001 | 0.054 |
| Adjusted | 364 | 0.3 (0.2, 0.4) | <0.001 | 0.237 | 0.3 (0.2, 0.4) | <0.001 | 0.217 | 0.3 (0.2, 0.4) | <0.001 | 0.247 |
| **CRP (mg/L)** |  |  |  |  |  |  |  |  |  |  |
| Unadjusted | 1087 | 0.8 (0.7, 1.0)<sup>a</sup> | <0.001 | 0.093 | 0.8 (0.6, 0.9)<sup>a</sup> | <0.001 | 0.126 | 1.3 (1.1, 1.4)<sup>b</sup> | <0.001 | 0.178 |
| Adjusted | 1039 | 0.8 (0.6, 1.0) | <0.001 | 0.191 | 0.9 (0.7, 1.0) | <0.001 | 0.205 | 0.9 (0.8, 1.1) | <0.001 | 0.221 |
| **Adiponectin (μg/mL)** |  |  |  |  |  |  |  |  |  |  |
| Unadjusted | 832 | -0.7 (-0.9, -0.4)<sup>a</sup> | <0.001 | 0.027 | -0.9 (-1.1, -0.6)<sup>a</sup> | 0.001 | 0.043 | -0.3 (-0.6, 0.1)<sup>b</sup> | 0.098 | 0.003 |
| Adjusted | 787 | -0.3(-0.6, -0.1) | 0.017 | 0.092 | -0.5 (-0.8, -0.2) | 0.001 | 0.099 | -0.2 (-0.5, 0.1) | 0.142 | 0.072 |
| **Total cholesterol (mmol/L)** |  |  |  |  |  |  |  |  |  |  |
| Unadjusted | 1440 | 0.1 (0.04, 0.1) | <0.001 | 0.009 | 0.1 (-0.003, 0.1) | 0.066 | 0.002 | 0.03 (-0.03, 0.1) | 0.323 | 0.001 |
| Adjusted | 1389 | 0.1 (0.04, 0.2) | <0.001 | 0.097 | 0.04 (-0.02, 0.1) | 0.127 | 0.090 | 0.04 (-0.01, 0.1) | 0.139 | 0.091 |
| **HDL cholesterol (mmol/L)** |  |  |  |  |  |  |  |  |  |  |
| Unadjusted | 1442 | -0.09 (-0.1, -0.1) | <0.001 | 0.066 | -0.1 (-0.1, -0.1) | <0.001 | 0.111 | -0.1 (-0.1, -0.04) | <0.001 | 0.030 |
| Adjusted | 1391 | -0.04 (-0.1, -0.03) | <0.001 | 0.216 | -0.1 (-0.1, -0.1) | <0.001 | 0.237 | -0.1 (-0.1, -0.03) | <0.001 | 0.221 |
| **LDL cholesterol (mmol/L)** |  |  |  |  |  |  |  |  |  |  |
| Unadjusted | 1436 | 0.1 (0.1, 0.2) | <0.001 | 0.018 | 0.09 (0.1, 0.1) | <0.001 | 0.012 | 0.06 (0.01, 0.1) | 0.014 | 0.004 |
| Adjusted | 1386 | 0.1 (0.1, 0.1) | <0.001 | 0.064 | 0.07 (0.02, 0.1) | 0.003 | 0.058 | 0.06 (0.01, 0.1) | 0.015 | 0.058 |
| **Triglycerides (mmol/L)** |  |  |  |  |  |  |  |  |  |  |
| Unadjusted | 1430 | 0.2 (0.1, 0.2) | <0.001 | 0.072 | 0.1 (0.2, 0.2) | <0.001 | 0.061 | 0.1 (0.1, 0.1) | <0.001 | 0.024 |
| Adjusted | 1380 | 0.1 (0.1, 0.2) | <0.001 | 0.185 | 0.1 (0.1, 0.1) | <0.001 | 0.173 | 0.1 (0.1, 0.1) | <0.001 | 0.166 |
| **FFA (mmol/L)** |  |  |  |  |  |  |  |  |  |  |
| Unadjusted | 1063 | 0.03 (0.02, 0.04) | <0.001 | 0.018 | 0.01 (0.00, 0.03) | 0.058 | 0.003 | 0.5 (0.04, 0.07) | <0.001 | 0.048 |
| Adjusted | 1022 | 0.04 (0.02, 0.05) | <0.001 | 0.203 | 0.04 (0.02, 0.05) | <0.001 | 0.189 | 0.04 (0.02, 0.05) | <0.001 | 0.199 |
| **Systolic blood pressure (mmHg)** |  |  |  |  |  |  |  |  |  |  |
| Unadjusted | 1499 | 3.8 (3.1, 4.5)<sup>a</sup> | <0.001 | 0.072 | 3.9 (3.2, 4.6)<sup>a</sup> | <0.001 | 0.073 | 1.9 (1.3, 2.6)<sup>b</sup> | <0.001 | 0.017 |
| Adjusted | 1445 | 2.9 (2.2, 3.6) | <0.001 | 0.209 | 2.5 (1.7, 3.2) | <0.001 | 0.199 | 2.6 (1.9, 3.4) | <0.001 | 0.203 |
| **Diastolic blood pressure (mmHg)** |  |  |  |  |  |  |  |  |  |  |
| Unadjusted | 1499 | 2.8 (2.8, 3.3)<sup>a</sup> | <0.001 | 0.068 | 2.1 (1.6, 2.6)<sup>a</sup> | <0.001 | 0.038 | 0.9 (0.4, 1.4)<sup>b</sup> | <0.001 | 0.007 |
| Adjusted | 1445 | 3.5 (2.9, 4.0) | <0.001 | 0.188 | 2.4 (1.8, 3.0) | <0.001 | 0.142 | 2.3 (1.8, 2.9) | <0.001 | 0.143 |

Data shown as estimated slope and confidence interval (CI) from multiple linear regression models. The adjusted models included age, gender, smoking and study.<sup>a,b</sup> Values with different superscripts letters indicate statistically significant differences between studies assessed by chi-squared test, $p < 0.05$ . Abbreviations: CRP, C-reactive protein; FFA, Free fatty acids; HOMA-IR, Homeostatic model assessment for insulin resistance; IL-6, Interleukin 6; SAD, sagittal abdominal diameter; TNF- $\alpha$ , Tumor necrosis factor alpha; WC, waist circumference.

### The ability of anthropometric measurements to identify individuals with metabolic syndrome markers above the IDF cut-offs

We found that BMI, WC and SAD were overall fair in their ability to identify individuals with metabolic syndrome markers above the IDF cut-offs after adjustments for age, sex, smoking and study and there was no differences between BMI, WC and SAD in their ability to identify these individuals (Table 3). However, in the unadjusted analysis WC and SAD performed slightly better than BMI in identifying individuals with metabolic syndrome markers above the IDF cut-offs for glucose >6.1 mmol/L, triglycerides, systolic and diastolic blood pressure, compared to BMI (Table 3). Using WC-to-height ratio and SAD-to-height ratio for identification of individuals with metabolic syndrome markers above the IDF cut-offs did not show any differences between the two (Supplementary table 2) or between these and SAD and WC alone.

**Table 3.** SAD, WC, and BMI as identifiers of individuals with metabolic syndrome markers above the IDF cut-offs measured as area under the receiver operating characteristics curve (AUROC) (n=1516).

|  | **SAD (cm)** | **WC (cm)** | **BMI (kg/m<sup>2</sup>)** |
| --- | --- | --- | --- |
| **Glucose (>5.6 mmol/L)** |  |  |  |
| Unadjusted | 0.53 (0.50, 0.57) | 0.56 (0.52, 0.59) | 0.53 (0.50, 0.56) |
| Adjusted | 0.79 (0.77, 0.82) | 0.80 (0.77, 0.83) | 0.80 (0.77, 0.82) |
| **Glucose (>6.1 mmol/L)** |  |  |  |
| Unadjusted | 0.60 (0.55, 0.64)<sup>a,b</sup> | 0.61 (0.57, 0.65)<sup>a</sup> | 0.52 (0.47, 0.56)<sup>b</sup> |
| Adjusted | 0.78 (0.74, 0.82) | 0.78 (0.75, 0.82) | 0.78 (0.74, 0.82) |
| **Triglycerides (>1.7 mmol/L)** |  |  |  |
| Unadjusted | 0.63 (0.60, 0.67)<sup>a</sup> | 0.62 (0.58, 0.65)<sup>a,b</sup> | 0.56 (0.52, 0.59)<sup>b</sup> |
| Adjusted | 0.72 (0.69, 0.76) | 0.71 (0.68, 0.74) | 0.71 (0.67, 0.74) |
| **HDL cholesterol (< 1.03 (men) or <1.29 (women) mmol/L)** |  |  |  |
| Unadjusted | 0.59 (0.56, 0.62) | 0.60 (0.57, 0.63) | 0.58 (0.55, 0.61) |
| Adjusted | 0.65 (0.62, 0.68) | 0.66 (0.63, 0.69) | 0.64 (0.61, 0.67) |
| **Systolic blood pressure (> 130 mmHg)** |  |  |  |
| Unadjusted | 0.63 (0.60, 0.66)<sup>a</sup> | 0.62 (0.59, 0.65)<sup>a</sup> | 0.55 (0.52, 0.58)<sup>b</sup> |
| Adjusted | 0.74 (0.72, 0.77) | 0.73 (0.71, 0.76) | 0.73 (0.71, 0.76) |
| **Diastolic blood pressure (>85 mmHg)** |  |  |  |
| Unadjusted | 0.62 (0.59, 0.65)<sup>a</sup> | 0.59 (0.56, 0.62)<sup>a,b</sup> | 0.53 (0.50, 0.57)<sup>b</sup> |
| Adjusted | 0.72 (0.69, 0.76) | 0.70 (0.67, 0.73) | 0.70 (0.67, 0.73) |

Data shown as estimated AUROC and 95% confidence interval using logistic regression models. The adjusted models included age, gender, smoking and study. Values with different superscripts letters indicate statistically significant differences between anthropometric measurements assessed by overlapping confidence intervals.

### Differences between sexes and between BMI categories

Contrary to the differences observed between the three anthropometric measures in the unadjusted models in men and women combined, there were no differences between BMI, WC and SAD in men or women alone (Supplementary table 3). However, we observed sex differences in the measures ability of the measures to identify individuals with metabolic syndrome markers above the IDF cut-offs. The prediction was generally better in women than in men especially for glucose >6.1 mmol/L, HDL and sBP in the unadjusted models and the prediction for sBP was still better for both SAD and BMI after adjustment (Supplementary table 3).

As for the models for the entire population, WC and SAD had a better ability to identify individuals with glucose >6.1 mmol/L, TAG, sBP, and dBP above the IDF cut-offs than BMI, but only in the unadjusted models and only in those with a BMI <35 kg/m^2^ (Supplementary table 4). For individuals with a BMI >35 kg/m^2^ a differences was only observed for glucose >5.6 mmol/L, where BMI and WC showed better associations that SAD. We found no differences in the associations at high and low BMI except for TAG, which had a higher AUROC at lower BMI even after adjustment. Similar tendencies were observed in the unadjusted models for all metabolic syndrome markers except HDL-cholesterol (Supplementary table 4).

## Discussion

In the present study, we found distinct differences between SAD and WC over the BMI range, which may indicate differences in adipose distribution across BMI categories. However, we did not find that any of the anthropometric measures were superior in the association with cardiometabolic health markers or in identifying people/subjects? with metabolic syndrome markers above the IDF cut-offs. Moreover, we did not find that one anthropometric measurement were superior depending on sex or BMI category. Finally, adjusting SAD and WC for height did not improve the identification ability of the anthropometric measures.

Earlier studies comparing SAD and WC show conflicting results and only examine a few risk markers at a time whereas our study measures multiple cardiometabolic risk markers giving a more complete metabolic overview. Similar to other studies our study show no difference between SAD and WC as markers of cardiometabolic health (19,20). One study in 826 elderly Dutch men and women report differences in correlations with markers of insulin resistance, dyslipidemia, and blood pressure between SAD or WC both above and below the age of 65 years of age but no consistently with respect to which was the better (19). However, other studies have shown that SAD is a superior anthropometric measure compared to WC and BMI (2,21,22) although some studies only show a minor benefit of SAD compared to WC (2,22). One study showed that SAD was a better predictor of insulin resistance compared to WC in high-risk group of 59 moderately obese men (21), whereas a study including 885 men and women only found that SAD was marginally better correlated with total and LDL-cholesterol, blood pressure and serum glucose and insulin (22). The latter is in line with results from a study with 4032 participants which showed small improvements in identifying elevated cardiometabolic risk in men with SAD compared to WC, but no difference women (2)

For hard endpoints such as CVD, SAD has been shown to be a better predictor than WC (4,9). One study showed that SAD >25 cm was the only anthropometric measurement associated with major CVD events in patients with T2D (4). Similary, a different study showed that SAD was the only measure that significantly predicted CVD in men, but not in women (9). This study also showed that SAD could predict CVD after adjustments for traditional biomarkers such as total- and HDL cholesterol and systolic blood pressure. This indicates that SAD might have predictive ability beyond established biomarkers of CVD. In the present study we tried to address this by including additional biomarkers linked to abdominal obesity, CVD and T2D including inflammatory cytokines. However, we did not find that SAD and WC were differentially associated with inflammatory markers.

The proposed mechanism behind the association of abdominal obesity and an adverse cardiometabolic profile is that increased adipose tissue mass and adipocyte size, leads to amplification in pro-inflammatory response of adipose tissue and raise FFA release into circulation (23,24). The increase of FFA and pro-inflammatory cytokines in circulation may increase the fat accumulation in skeletal muscle and liver which may lead to insulin resistance and related cardiometabolic abnormalities (23,24). Especially VAT is associated with increased cytokine production and FFA release into circulation. In the present study, we included inflammatory cytokines in order to elucidate whether SAD was really, a better predictor of adverse metabolic cytokine production associated with increased VAT. We did not find that SAD and WC were differentially associated with inflammatory markers and thus cannot shed any light on whether this could explain the before mentioned superior predictive ability of SAD on top of established risk markers.

The discrepancies in results between studies might also be because of age, ethnicity, sex, phenotype or methodological differences. One of the methodological differences that might lead to discrepancies in the results is the choice measurement sites and the optimal point of measuring SAD is still debated (3). We used the highest point of the abdomen, which differ from the protocol described in the US National Health and Nutrition Examination Survey (NHANES). This might result in lower correlation with cardiometabolic measures (3,25,26) and could prevent us from observing differences between SAD and WC. However, if minor differences in measurement point can affect the associations drastically, its clinical utility may be limited. Furthermore, it has been argued that assessment of WC requires less tools and is more practical compared to SAD, which might limit the use of SAD as a clinical screening tool. However, similarly to others(2) we find that difference is negligible. SAD has also shown to have a high intraclass correlation in both lean and obese individuals, while the intraclass correlation was lower in obese compared to lean individuals when measuring WC (27). This could indicate that SAD is more reliable when measuring individuals of different weight categories and that SAD is better than WC in individuals with high amounts of abdominal fat, especially SAT. SAD may therefore be more reliable when measuring individuals of different weight categories with low variation and might also be less sensitive to the investigator performing the measurement compared to WC.

One limitation of the present study is the cross-sectional design using biomarkers which is a weaker design compared to studies with a prospective design and hard endpoints such as T2D, CVD and mortality. Such studies are highly warranted. Furthermore, the present study used individuals, which were primarily overweight and obese (98%), whereas other studies that used both normal weight and overweight individuals (7,8). This limits our generalizability to normal weight populations. However, our study used a clinically relevant population group, which are at risk of developing T2D and CVD, and thus a clear relevance for establishing a highly predictable anthropometric measure. The current study has measured many different cardiometabolic outcomes including markers of insulin resistance, dyslipidemia, low grade inflammation and blood pressure. As previous studies have not included all these markers, this study provides a good view of overall cardiometabolic health. Furthermore, the sample size in the present study is appropriate to examine the research questions posed.

## Conclusions

In conclusion, the data from 6 cohorts of high-risk individuals showed that neither SAD, WC nor BMI seems to be superior as an indicator of an adverse metabolic phenotype or in their ability to identify individuals with metabolic syndrome markers above the IDF cut-offs regardsless of sex and BMI category. Thus choosing the most practical anthropometric measure might be the best solution from a public health perspective.

## Abbreviations

BMI: body mass index
CT: computed tomography
FFA: free fatty acids
IDF: International Diabetes Federation
IL: interleukin
MRI: magnetic resonance imaging
ROC: Receiver Operating Characteristic
SAD: sagittal abdominal diameter
SAT: subcutaneous adipose tissue
TNF: tumor necrosis factor
VAT: visceral adipose tissue
WC: waist circumference.

## Acknowledgements

The authors thank all the study participants and the staff who have contributed to the planning and conduction of the studies.

## Author contributions

MVL, MK, AA, TML, CR and GM were involved in conception and design of the study. MVL, LK, SV, SKK, OP, LL, TML, WS, AA and MK were involved in the collection of data and biological samples. CR, GM and MVL did the statistical analysis. MVL and GM drafted the manuscript. All authors were involved in the interpretation of the data and all authors read, revised and approved the final manuscript.

