## Supplemental table 1 for "Sagittal abdominal diameter and waist circumference are equally good as identifiers of cardiometabolic risk"

**Supplementary method section**

**Anthropometric**

The methodology of the anthropometric measurements has been described in detail elsewhere (1–5). In brief, all participants voided before anthropometric measurements were performed. Body weight was measured with individuals dressed in light clothing or underwear and without shoes. Body weight was noted to the nearest 0.05 kg. Height was measured twice with 0.5 cm accuracy without shoes using a wall-mounted stadiometer and the average of the two measurements was recorded: BMI was calculated as weight (kg) divided by the square of height (m^2^). After 10 min rest, BP was measured using a digital blood pressure monitor with participants in a vertical position after at least 10 minutes rest. Blood pressure was measured three consecutive times and a mean value of the three measurements was used in subsequent analyses.

**Biochemical analyses of blood samples**

The methodology of the biochemical analyses has been described in detail elsewhere (3–8). In brief, venous blood was drawn from antecubital vein after overnight fasting. In 3G, RIGHT and PROKA, glucose was analysed by an enzymatic method using Hexokinase and Glucose-6-phosphate-dehydrogenase. The analysis was carried out by using an ABX Pentra 400 chemistry analyzer (ABX Pentra, Horiba ABX, Montpellier, France). Insulin was measured by a chemiluminescent immunometric assay (Immulite 1000, Siemens Medical Solutions Diagnostics, Los Angeles, USA) (3,5,6). In Diogenes, fasting blood samples were drawn from a venflon and were followed by an oral glucose tolerance test (OGTT). The individuals consumed 75 g of glucose diluted with 250 ml of water. Blood samples were drawn multiple times during the 120 min test time. Fasting and OGTT serum glucose were analyzed by colorimetric assays (Ortho-Clinical Diagnostics, Johnson & Johnson, Birkerød, Denmark). Fasting insulin concentrations were measured by a solid-phase, two-site chemiluminescent immunometric assay (Siemens Medical Solutions Diagnostics, Ballerup, Denmark) for the IMMULITE 2500 analyzer (7). In OPUS, fasting blood samples were obtained from an intravenous catheter in the antecubital vein. Glucose was analyzed using Vitros reagents on a Vitros 5.1 FS (Ortho-clinical diagnostics, Johnson & Johnson, Denmark). Blood samples were followed by a 120 min OGTT similar to the Diogenes study. Serum insulin was analyzed using a solid-phase, sandwich chemiluminescent immunometric assay on ADVIA Centauer XP (Siemens, Denmark) (4).

Based on the measured values of fasting glucose and insulin, the homeostatic model assessment for insulin resistance (HOMA-IR) was calculated according to the following equation: fasting glucose (mmol/L) * fasting insulin (mU/L))/22.5 (9). Based on the measured values of glucose and insulin during the OGTT the Matsuda index was calculated according to the following equation: 10,000/$\surd$ ((fasting plasma glucose (mg/dl) * fasting plasma insulin (mU/ml)) * (mean OGTT glucose (mg/dl) * mean OGTT insulin (mU/ml))) (10).

In 3G and RIGHT, Serum inflammatory markers IL-6, IL1β and TNF-α were measured using high-sensitivity enzyme linked immunosorbent assay (ELISA) (HS600B and HSTA00D, R&D systems, Minneapolis, Minnesota, USA) (3,6). In Diogenes, serum hsCRP was measured via an immunoturbidimetric assay (Roche Diagnostics), with use of monoclonal antibodies to CRP and a colorimetric assay (Roche Diagnostics), for the COBAS Integra 400 analyzer (7). In OPUS, CRP were analyzed using Vitros reagents on a Vitros 5.1 FS (Ortho-clinical diagnostics, Johnson & Johnson, Denmark) and TNF-α and adiponectin concentration in plasma were measured by ELISA (R&D Systems, Minneapolis, USA) (4). For all studies included in the present study, values below the lower limit of quantification for CRP were set to 0.001 indicating very low concentrations (1,3–5,7,8). For all six intervention trials, analysis was carried out excluding CRP values >10 mg/L, indicating acute illness and not low grade inflammation. In 3G and RIGHT, total-, LDL- and HDL-cholesterol, triglycerides, and FFA were analysed using automated, enzymatic, colorimetric assay on ABX 4 Pentra 400 chemistry analyzer (ABX Pentra, Horiba ABX, Montpellier, France) (3,6). In Diogenes, fasting serum total cholesterol, HDL cholesterol and triglycerides were measured by routine enzymatic assays (Roche Diagnostics, Hvidovre, Denmark) in the COBAS Integra 400 analyzer. In OPUS, total cholesterol, HDL and triglycerides were analyzed using Vitros reagents on a Vitros 5.1 FS (Ortho-clinical diagnostics, Johnson & Johnson, Denmark). In both studies, low-density lipoprotein (LDL) cholesterol was calculated from total cholesterol, HDL and triglycerides by the Friedewald equation (7). In PROKA, total cholesterol and triglycerides were measured via enzymatic photometric tests and HDL, LDL and FFA were measured via enzymatic colorimetric tests (5).

**Supplemental table 1** Associations between baseline SAD/height ratio and WC/height ratio and baseline markers

|  |  | **SAD/height ratio** | | | **WC/height ratio** | | |
| --- | --- | --- | --- | --- | --- | --- | --- |
|  | **n** | **Slope (CI)** | ***p*-value** | **R^2^** | **Slope (CI)** | ***p*-value** | **R^2^** |
| **Fasting insulin** (pmol/L) | | | | | | | |
| Unadjusted | 1405 | 12.9 (11.0, 14.9) | <0.001 | 0.109 | 15.1 (13.2, 17.0) | <0.001 | 0.145 |
| Adjusted | 1354 | 13.7 (11.5, 15.8) | <0.001 | 0.164 | 16.4 (14.3, 18.5) | <0.001 | 0.202 |
| **Fasting glucose** (mmol/L) | | | | | | | |
| Unadjusted | 1426 | 0.01 (-0.03, 0.1) | 0.580 | 0.000 | 0.04 (0.01, 0.1) | 0.015 | 0.004 |
| Adjusted | 1376 | 0.1 (0.1, 0.1) | <0.001 | 0.252 | 0.1 (0.1, 0.2) | <0.001 | 0.263 |
| **2-h glucose** (mmol/L) | | | | | | | |
| Unadjusted | 975 | 0.4 (0.3, 0.5)^a^ | <0.001 | 0.043 | -0.5 (-0.8, -0.2)^b^ | 0.000 | 0.013 |
| Adjusted | 925 | 0.3(0.2, 0.5) | <0.001 | 0.093 | 0.3 (0.2, 0.4) | <0.001 | 0.090 |
| **HOMA-IR** | | | | | | | |
| Unadjusted | 1404 | 0.5 (0.5, 0.6) | <0.001 | 0.122 | 0.6 (0.6, 0.7) | <0.001 | 0.165 |
| Adjusted | 1353 | 0.5 (0.4, 0.6) | <0.001 | 0.242 | 0.6 (0.5, 0.7) | <0.001 | 0.280 |
| **Matsuda index** | | | | | | | |
| Unadjusted | 963 | -0.9 (-1.0, -1.0) | <0.001 | 0.102 | -1.1 (-1.2, -1.0) | <0.001 | 0.161 |
| Adjusted | 913 | -0.9 (-1.0, -1.0) | <0.001 | 0.149 | -1.1 (-1.3, -1.0) | <0.001 | 0.208 |
| **TNF-α** (pg/mL) | | | | | | | |
| Unadjusted | 298 | 0.1 (0.02, 0.2) | 0.018 | 0.019 | 0.1 (0.02, 0.2) | 0.011 | 0.022 |
| Adjusted | 292 | 0.1 (0.02, 0.2) | 0.016 | 0.226 | 0.1 (0.02, 0.2) | 0.014 | 0.226 |
| **IL-6** (pg/mL) | | | | | | | |
| Unadjusted | 369 | 0.3 (0.2, 0.4) | <0.001 | 0.068 | 0.2 (0.1, 0.3) | <0.001 | 0.061 |
| Adjusted | 364 | 0.3 (0.2, 0.4) | <0.001 | 0.240 | 0.3 (0.2, 0.4) | <0.001 | 0.225 |
| **CRP** (mg/L) | | | | | | | |
| Unadjusted | 1087 | 0.8 (0.7, 1.0) | <0.001 | 0.139 | 0.9 (0.8, 1.0) | 0.000 | 0.151 |
| Adjusted | 1039 | 0.8 (0.7, 0.9) | <0.001 | 0.201 | 0.9 (0.8, 1.03) | <0.001 | 0.219 |
| **Adiponectin** (µg/mL) | | | | | | | |
| Unadjusted | 832 | -0.4 (-0.6, -0.1) | 0.013 | 0.007 | -0.5 (-0.8, -0.2) | 0.002 | 0.012 |
| Adjusted | 787 | -0.3 (-0.5, 0.03) | 0.080 | 0.089 | -0.4 (-0.7, -0.1) | 0.001 | 0.093 |
| **Total cholesterol** (mmol/L) | | | | | | | |
| Unadjusted | 1440 | 0.1 (0.1, 0.2) | <0.001 | 0.017 | 0.1 (0.04, 0.1) | <0.001 | 0.009 |
| Adjusted | 1389 | 0.1 (0.1, 0.2) | <0.001 | 0.107 | 0.1 (0.04,0.1) | <0.01 | 0.096 |
| **HDL cholesterol** (mmol/L) | | | | | | | |
| Unadjusted | 1442 | -0.1 (-0.1, -0.04) | <0.001 | 0.026 | -0.1 (-0.1, -0.1) | <0.001 | 0.050 |
| Adjusted | 1391 | -0.04 (-0.1, -0.02) | <0.001 | 0.213 | -0.1 (-0.1, -0.04) | <0.001 | 0.230 |
| **LDL cholesterol** (mmol/L) | | | | | | | |
| Unadjusted | 1436 | 0.1 (0.1, 0.2) | <0.001 | 0.025 | 0.1 (0.1, 0.2) | <0.001 | 0.020 |
| Adjusted | 1386 | 0.1 (0.1, 0.2) | <0.001 | 0.074 | 0.1 (0.1, 0.2) | <0.001 | 0.068 |
| **Triglycerides** (mmol/L) | | | | | | | |
| Unadjusted | 1430 | 0.1 (0.1, 0.2) | <0.001 | 0.052 | 0.1 (0.1, 0.2) | <0.001 | 0.040 |
| Adjusted | 1380 | 0.1 (0.1, 0.2) | <0.001 | 0.190 | 0.1 (0.1, 0.1) | <0.001 | 0.178 |
| **FFA** (mmol/L) | | | | | | | |
| Unadjusted | 1063 | 0.1 (0.04, 0.1) | <0.001 | 0.047 | 0.04 (0.02, 0.1) | <0.001 | 0.027 |
| Adjusted | 1022 | 0.04 (0.03, 0.1) | <0.001 | 0.205 | 0.03 (0.01, 0.04) | <0.001 | 0.191 |
| **Systolic blood** **pressure** (mmHg) | | | | | | | |
| Unadjusted | 1499 | 2.7 (2.0, 3.4) | <0.001 | 0.036 | 2.5 (1.8, 3.2) | <0.001 | 0.031 |
| Adjusted | 1445 | 2.6 (1.9, 3.3) | <0.001 | 0.205 | 2.1 (1.4, 2.8) | <0.001 | 0.195 |
| **Diastolic blood pressure** (mmHg) | | | | | | | |
| Unadjusted | 1499 | 2.1 (1.6, 2.7) | <0.001 | 0.039 | 1.3 (0.7, 1.8) | <0.001 | 0.014 |
| Adjusted | 1445 | 3.1 (2.5, 3.6)^a^ | <0.001 | 0.174 | 1.9 (1.3, 2.5)^b^ | <0.001 | 0.132 |

Data are shown as estimated slope and confidence interval (CI) from multiple linear regression models. The adjusted models included controlling for age, gender, smoking and study. Values with different superscripts letters indicate statistically significant differences between anthropometric measurements assessed by overlapping confidence intervals, p< 0.05. Abbreviations: CRP, C-reactive protein; FFA, Free fatty acids; HOMA-IR, Homeostatic model assessment for insulin resistance; IL-6, Interleukin 6, SAD, sagittal abdominal diameter; TNF- α, Tumor necrosis factor alpha; WC, waist circumference.

**Supplemental table 2** SAD/height ratio and WC/height ratio as identifiers of individuals with metabolic syndrome markers above the IDF cut-offs measured as area under the receiver operating characteristics (ROC) curve.

|  | **n** | **SAD/height ratio** | **WC/height ratio** |
| --- | --- | --- | --- |
| **Glucose** (>5.6 mmol/L) | | | |
| Unadjusted | 1516 | 0.51 (0.48, 0.54) | 0.48 (0.44, 0.51) |
| Adjusted | 1516 | 0.79 (0.77, 0.82) | 0.80 (0.77, 0.83) |
| **Glucose** (>6.1 mmol/L) | | | |
| Unadjusted | 1516 | 0.57 (0.53, 0.62) | 0.58 (0.54, 0.63) |
| Adjusted | 1516 | 0.78 (0.74, 0.82) | 0.79 (0.75, 0.82) |
| **Triglycerides** (>1.7 mmol/L) | | | |
| Unadjusted | 1516 | 0.60 (0.57, 0.63) | 0.58 (0.54, 0.61) |
| Adjusted | 1516 | 0.72 (0.69, 0.75) | 0.71 (0.68, 0.74) |
| **HDL cholesterol** (< 1.03 (men) or <1.29 (women) mmol/L) | | | |
| Unadjusted | 1516 | 0.58 (0.55, 0.61) | 0.52 (0.56, 0.62) |
| Adjusted | 1516 | 0.64 (0.61, 0.67) | 0.65 (0.62, 0.68) |
| **Systolic blood** **pressure** (>130 mmHg) | | | |
| Unadjusted | 1516 | 0.59 (0.56, 0.62) | 0.58 (0.55, 0.61) |
| Adjusted | 1516 | 0.74 (0.72, 0.77) | 0.73 (0.70, 0.76) |
| **Diastolic blood pressure** (> 85 mmHg) | | | |
| Unadjusted | 1516 | 0.60 (0.56, 0.63) | 0.56 (0.53, 0.59) |
| Adjusted | 1516 | 0.72 (0.69, 0.75)^a^ | 0.65 (0.64, 0.68)^b^ |

Data are shown as estimated area under the curve and 95% confidence interval (CI) for the receiver operating characteristics (ROC) curve using logistic regression models. The adjusted models included controlling for age, gender, smoking and study.

**Supplemental table 3** SAD, WC, and BMI as identifiers of individuals with metabolic syndrome markers above the IDF cut-offs measured as area under the receiver operating characteristics (ROC) curve.

|  | **Men** | | | | **Women** | | | |
| --- | --- | --- | --- | --- | --- | --- | --- | --- |
|  | **n** | **SAD** (cm) | **WC** (cm) | **BMI** (kg/m^2^) | **n** | **SAD** (cm) | **WC** (cm) | **BMI** (kg/m^2^) |
| **Glucose** (> 5.6 mmol/L) | | | | | | | | |
| Unadjusted | 485 | 0.55 (0.50, 0.61) | 0.55 (0.49, 0.60) | 0.59 (0.54, 0.65) | 1031 | 0.55 (0.51, 0.59) | 0.56 (0.52, 0.61) | 0.51 (0.47, 0.55) |
| Adjusted | 485 | 0.80 (0.76, 0.85) | 0.81 (0.76, 0.85) | 0.81 (0.76, 0.85) | 1031 | 0.79 (0.75, 0.82) | 0.80 (0.76, 0.83) | 0.78 (0.75, 0.82) |
| **Glucose** (> 6.1 mmol/L) | | | | | | | | |
| Unadjusted | 485 | 0.49 (0.42, 0.57) | 0.50 (0.43, 0.58) | 0.55 (0.48, 0.63) | 1031 | 0.64 (0.58, 0.69) | 0.64 (0.59, 0.70) | 0.57 (0.51, 0.63) |
| Adjusted | 485 | 0.75 (0.69, 0.82) | 0.76 (0.69, 0.82) | 0.76 (0.70, 0.82) | 1031 | 0.79 (0.75, 0.84) | 0.80 (0.75, 0.84) | 0.78 (0.73, 0.83) |
| **Triglycerides** (>1.7 mmol/L) | | | | | | | | |
| Unadjusted | 485 | 0.57 (0.52, 0.62) | 0.54 (0.49, 0.60) | 0.56 (0.51, 0.61) | 1031 | 0.62 (0.58, 0.67) | 0.59 (0.54, 0.63) | 0.56 (0.51, 0.61) |
| Adjusted | 485 | 0.64 (0.59, 0.70) | 0.64 (0.59, 0.70) | 0.65 (0.59, 0.70) | 1031 | 0.70 (0.66, 0.75) | 0.68 (0.63, 0.72) | 0.65 (0.60, 0.70) |
| **HDL cholesterol** (< 1.03 (men) or <1.29 (women) mmol/L) | | | | | | | | |
| Unadjusted | 485 | 0.54 (0.49, 0.59) | 0.53 (0.48, 0.58) | 0.54 (0.48, 0.59) | 1031 | 0.63 (0.59, 0.66) | 0.64 (0.61, 0.68) | 0.61 (0.57, 0.64) |
| Adjusted | 485 | 0.64 (0.59, 0.69) | 0.64 (0.59, 0.69) | 0.65 (0.60, 0.70) | 1031 | 0.67 (0.64, 0.71) | 0.69 (0.65, 0.72) | 0.66 (0.62, 0.69) |
| **Systolic blood** **pressure** (> 130 mmHg) | | | | | | | | |
| Unadjusted | 485 | 0.55 (0.49, 0.58) | 0.53 (0.48, 0.58) | 0.51 (0.46, 0.56) | 1031 | 0.63 (0.59, 0.67) | 0.60 (0.56, 0.64) | 0.58 (0.54, 0.62) |
| Adjusted | 485 | 0.63 (0.58, 0.68) | 0.62 (0.57, 0.67) | 0.62 (0.57, 0.67) | 1031 | 0.74 (0.70, 0.77) | 0.72 (0.68, 0.76) | 0.72 (0.69,0.76) |
| **Diastolic blood pressure** (> 85 mmHg) | | | | | | | | |
| Unadjusted | 485 | 0.60 (0.55, 0.66) | 0.55 (0.50, 0.60) | 0.53 (0.47, 0.58) | 1031 | 0.60 (0.56, 0.65) | 0.57 (0.53, 0.61) | 0.54 (0.50, 0.58) |
| Adjusted | 485 | 0.69 (0.64, 0.74) | 0.66 (0.61, 0.71) | 0.66 (0.60, 0.71) | 1031 | 0.72 (0.68, 0.76) | 0.69 (0.65, 0.73) | 0.69 (0.66, 0.73) |

Data are shown as estimated area under the curve and 95% confidence interval (CI) for the receiver operating characteristics (ROC) curve using logistic regression models. The adjusted models included controlling for age, gender, smoking and study.

|  | **BMI ≤35** | | | | **BMI > 35** | | | | |
| --- | --- | --- | --- | --- | --- | --- | --- | --- | --- |
|  | **n** | **SAD** (cm) | **WC** (cm) | **BMI** (kg/m^2^) | **n** | **SAD** (cm) | **WC** (cm) | **BMI** (kg/m^2^) | |
| **Glucose** (> 5.6 mmol/L) | | | | | | | | | |
| Unadjusted | 1051 | 0.57 (0.53, 0.61) | 0.57 (0.53, 0.61) | 0.53 (0.49, 0.57) | 463 | 0.52(0.46, 0.58)^b^ | 0.58 (0.53, 0.64)^a^ | | 0.56 (0.49, 0.62)^a^ |
| Adjusted | 1051 | 0.81 (0.78, 0.84) | 0.81 (0.78, 0.84) | 0.81 (0.78, 0.84) | 463 | 0.77 (0.72, 0.82) | 0.78 (0.73, 0.83) | | 0.77 (0.71, 0.82) |
| **Glucose** (> 6.1 mmol/L) | | | | | | | | | |
| Unadjusted | 1051 | 0.63 (0.58, 0.68)^a^ | 0.63 (0.58, 0.68)^a^ | 0.52 (0.46, 0.57)^b^ | 463 | 0.58 (0.50;0.69) | 0.60 (0.52, 0.69) | | 0.52 (0.42, 0.61) |
| Adjusted | 1051 | 0.82 (0.77, 0.86) | 0.82 (0.77;0.86) | 0.82 (0.78, 0.86) | 463 | 0.71 (0.63, 0.81) | 0.74 (0.66, 0.81) | | 0.71 (0.63, 0.79) |
| **Triglycerides** (1.7 mmol/L) | | | | | | | | | |
| Unadjusted | 1051 | 0.66 (0.62, 0.70)^a^ | 0.66 (0.62, 0.70)^a^ | 0.57 (0.53, 0.61)^b^ | 463 | 0.62 (0.56, 0.68) | 0.60 (0.54, 0.66) | | 0.50 (0.44, 0.57) |
| Adjusted | 1051 | 0.75 (0.72, 0.79) | 0.74 (0.70, 0.78) | 0.73 (0.69, 0.77) | 463 | 0.67 (0.61, 0.73) | 0.67 (0.61, 0.73) | | 0.66 (0.60, 0.72) |
| **HDL cholesterol** (< 1.03 (men) or <1.29 (women) mmol/L) | | | | | | | | | |
| Unadjusted | 1051 | 0.59 (0.55, 0.62) | 0.59 (0.55, 0.62) | 0.57 (0.54, 0.61) | 463 | 0.54 (0.48, 0.59) | 0.55 (0.49, 0.60) | | 0.53 (0.48, 0.59) |
| Adjusted | 1051 | 0.65 (0.61, 0.68) | 0.66 (0.63, 0.70) | 0.64 (0.61, 0.68) | 463 | 0.68 (0.63, 0.73) | 0.69 (0.64, 0.74) | | 0.67 (0.62, 0.72) |
| **Systolic blood** **pressure** >130 mmHg) | | | | | | | | | |
| Unadjusted | 1051 | 0.64 (0.61, 0.68)^a^ | 0.64 (0.61, 0.68)^a^ | 0.54 (0.50, 0.58)^b^ | 463 | 0.65 (0.59, 0.70) | 0.62 (0.57, 0.68) | | 0.58 (0.53, 0.64) |
| Adjusted | 1051 | 0.76 (0.73, 0.79) | 0.75 (0.72, 0.78) | 0.75 (0.72, 0.78) | 463 | 0.72 (0.67, 0.76) | 0.70 (0.65, 0.75) | | 0.70 (0.65, 0.75) |
| **Diastolic blood pressure** (>85 mmHg) | | | | | | | | | |
| Unadjusted | 1051 | 0.61 (0.57, 0.65)^a^ | 0.61 (0.57, 0.65)^a^ | 0.51 (0.47, 0.55)^b^ | 463 | 0.68 (0.62, 0.73) | 0.61 (0.55, 0.67) | | 0.56 (0.50, 0.62) |
| Adjusted | 1051 | 0.74 (0.70, 0.77) | 0.72 (0.68, 0.75) | 0.71 (0.68, 0.75) | 463 | 0.71 (0.66, 0.76) | 0.66 (0.60, 0.72) | | 0.68 (0.62, 0.73) |

**Supplemental table 4** SAD, WC, and BMI as identifiers of individuals with metabolic syndrome markers above the IDF cut-offs measured as area under the receiver operating characteristics (ROC) curve.

Data are shown as estimated area under the curve and 95% confidence interval (CI) for the receiver operating characteristics (ROC) curve using logistic regression models. The adjusted models included controlling for age, gender, smoking and study. Values with different superscripts letters indicate statistically significant differences between anthropometric measurements assessed by overlapping confidence intervals.
